# Autonomous Shaping of the piRNA Sequence Repertoire by Competition between Adjacent Ping-Pong Sites

**DOI:** 10.1101/2024.05.22.595280

**Authors:** Jie Yu, Natsuko Izumi, Yukihide Tomari, Keisuke Shoji

## Abstract

PIWI-interacting RNAs (piRNAs) are crucial for silencing transposable elements (TEs). In many species, piRNAs are generated via a complex process known as the ping-pong pathway, which couples TE cleavage with new piRNA amplification. However, the biological significance of this complexity and its impact on the piRNA sequence repertoire remain unclear. Here, we systematically compared piRNA production patterns in two closely related silkworm cell lines and found significant changes in their piRNA sequence repertoire. Importantly, the changeability of this repertoire showed a strong negative correlation with the efficiency of piRNA biogenesis. This can be explained by competition between adjacent ping-pong sites, as supported by our mathematical modeling. Moreover, this competition can rationalize how piRNAs autonomously avoid deleterious mismatches to target TEs. These findings unveil the intrinsic plasticity and adaptability of the piRNA system to combat diverse TE sequences and highlight the universal power of competition and self-amplification to drive autonomous optimization.

## INTRODUCTION

Transposable elements (TEs) are mobile genetic elements that can cause mutations and disrupt the function of genes, and regulating TE activity is essential for organisms to maintain their genome integrity and stability^1,2^. piRNAs, a class of small non-coding RNAs ranging from 24 to 32 nucleotides (nt) in length, play a significant role in this regulation process in the animal germline^3-5^. piRNAs guide PIWI proteins to complementary sequences, silencing TEs at the transcriptional and/or post-transcriptional levels^6,7^.

The biogenesis of piRNAs starts from the transcription of piRNA precursors from TEs or genomic regions called piRNA clusters, which contain fragments of TEs^8,9^. piRNA precursors are then processed into mature piRNAs by two distinct mechanisms: the ping-pong pathway and the phased pathway^5^. In the ping-pong pathway, a PIWI protein guided by an initiator piRNA cleaves a complementary precursor transcript, yielding a 5’-monophosphorylated fragment called pre-pre-piRNA^9-12^. The 5’ portion of pre-pre-piRNA is then loaded into another PIWI protein, while its 3’ portion is cleaved off by the endonuclease Zucchini^13-15^ or at a downstream piRNA target site^12,15^ to produce pre-piRNA. The PIWI-loaded pre-piRNA is then further trimmed at its 3’ end by the exonuclease Trimmer to the mature length, producing a responder piRNA that overlaps with the original initiator piRNA by 10 nt at its 5’ ends^16^. In this way, the ping-pong pathway amplifies a sense:antisense pair of 10-nt overlapping piRNAs^17^. In the phased piRNA biogenesis pathway, the 3’ fragment resulting from Zucchini-mediated cleavage of a pre-pre-piRNA (the other fragment not used for the ping-pong pathway) is loaded into the next PIWI protein as a new pre-pre-piRNA^12^. This pre-pre-piRNA is once again cleaved by Zucchini, generating a pre-piRNA^13,14^, which is subsequently trimmed by Trimmer to produce a mature trailing piRNA. Accordingly, the phased pathway expands the repertoire of piRNAs in the 3’ direction^13,14^.

The ping-pong pathway plays a dual role: it silences TEs by cleavage while amplifying new piRNAs using the cleavage fragments^5,12,18^. Amplified piRNAs can then be maternally inherited by the germline of the next generation, serving as a robust trans-generational repository of TE sequence information for target silencing^19,20^. Indeed, if a TE, whose sequence is absent from the maternal pool of piRNAs, is inherited only paternally, the TE escapes silencing and causes adult sterility syndrome^6,21^. A classic example is P-M hybrid dysgenesis in *Drosophila*, where crosses between a male fly with *P* elements (P strain) and a female without *P* elements (M strain) cause sterility in the offspring, while reciprocal crosses between a P strain female and an M strain male produce fertile progeny because of maternally inherited piRNAs that target *P* elements^6,22^. In addition, piRNAs can tolerate more mismatches than small interfering RNAs (siRNAs), another class of small RNAs that guide AGO proteins, for target recognition and cleavage, making them suitable for combating rapidly diverging or newly invading TEs^20,23^. Still, efficient cleavage by piRNAs requires more than 15 contiguous base pairs^20^, which is expected to be particularly important in the ping-pong pathway for reciprocal cleavage cycles. Therefore, it must be critical for the piRNA system to choose the appropriate sites for ping-pong-mediated piRNA production and to have the flexibility to adjust them according to diverging TEs for achieving long-term TE silencing over generations.

Then, how are piRNA-producing sites determined? A subset of piRNAs shows a strong preference for uridine (U) at their 5’ end (called 1U bias)^24^, which can be explained by the structural feature of the PIWI proteins to which they bind^12,25^. However, not all Us within precursor transcripts are actually used for the production of the 5’ end of piRNAs, and it remains unknown how the piRNA-producing sites are selected and how flexible that selection can be. In this study, we conducted a comparative analysis of the small RNA sequencing data derived from two closely related silkworm cultured cells that were separated by continuous cultivation for seven years. We found that, although the total amounts of piRNAs per TE did not change significantly, some TEs drastically changed the positions of piRNA-producing sites. Importantly, TEs with less efficient piRNA biogenesis were more prone to changes in piRNA production patterns, and conversely, TEs with more efficient piRNA biogenesis showed more stable piRNA production patterns. This can be explained by competition between adjacent ping-pong sites, as supported by our mathematical modeling. Moreover, this competition model can rationalize how piRNAs, as a system, can organize themselves to avoid deleterious mismatches to target TEs. We propose that the intrinsic flexible nature of the piRNA system allows it to autonomously search for optimal patterns of piRNA production to combat rapidly diverging or newly acquired TEs.

## RESULTS

### piRNA production patterns are changeable over time

When analyzing our previous small RNA sequencing data derived from silkworm BmN4 cells^15,26^, we noticed that some but not all piRNAs show different production patterns depending on when the libraries were constructed. To gain a precise understanding of this observation, we decided to conduct a comparative analysis of small RNA sequencing data from two sibling BmN4 cell lines that were 7 years apart; one cell line, Naive20, was continuously cultured from 2013 to 2020, while the other cell line, Naive13, was stored in a deep freezer during that entire period (Figure 1A). To avoid potential variations across different batches of sequencing, we created the small RNA libraries of these two different types of cells at the same time and performed sequencing on the same flow cell. We also included two additional cell types where Naive13 or Naive20 cells were transiently transfected with dsRNAs against Renilla luciferase as mock RNAi, which in theory should not affect the gene expression profiles (dsRluc13 and dsRluc20; Figure S1A). We then mapped the piRNA reads to 1,811 TEs that are well defined in the silkworm genome^27^ and plotted the change in piRNA abundance per TE between the different libraries (Figures S1B–E).

**Figure 1.**
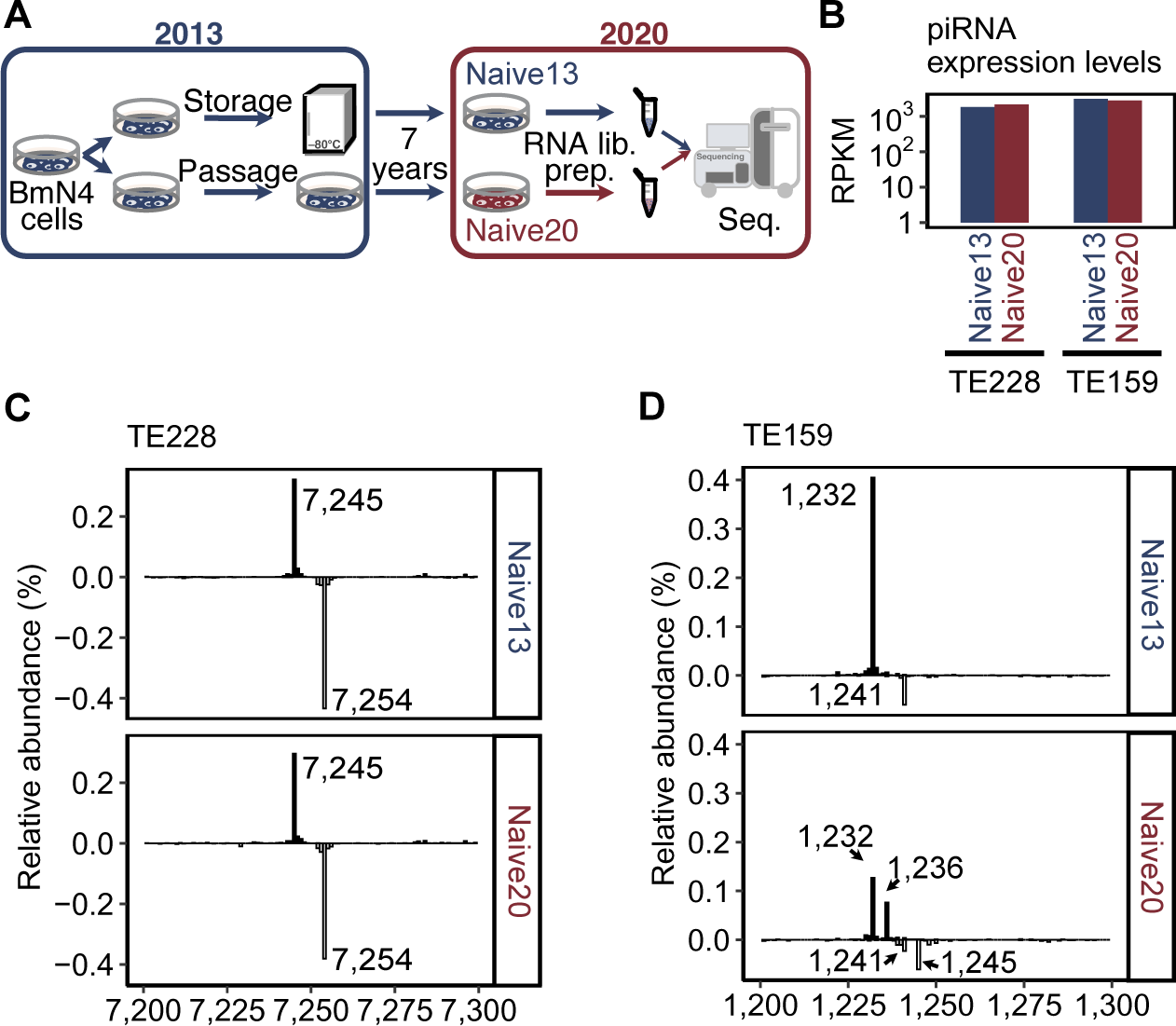
piRNA production patterns are changeable over time. (A) Scheme for the two cell lines used in this study. Naive20 was continuously cultured from 2013 to 2020, while Naive13 was stored at –80°C during that period. The small RNA libraries were prepared at the same time and sequenced on the same flow cell. (B) Bar graph showing piRNA abundances mapped to two representative TEs, TE228 (left) and TE159 (right), in Naive13 (dark blue) and Naive20 (dark red), respectively. (C) Distribution of the 5’ ends of piRNAs that mapped to a specific region of TE228, showing two main peaks at position 7,245 and position 7,254 in both Naive13 (top) and Naive20 (bottom). Total reads mapped to TE228 were used for normalization. (D) Distribution of the 5’ ends of piRNAs that mapped to a specific region of TE159, showing two main peaks at position 1,232 and position 1,241 in both Naive13 (top) and Naive20 (bottom), and two additional peaks at position 1,236 and position 1,245 in Naive20 (bottom). Total reads mapped to TE159 were used for normalization.

As expected, the abundance of TE-mapped piRNAs was almost identical between the two cell lines from the same period with or without mock RNAi, i.e., between Naive13 vs. dsRluc13 and Naive20 vs. dsRluc20 (Figures S1B and S1C). Even when cell lines from different periods were compared (Naive13 vs. Naive20 and dsRluc13 vs. dsRluc20), many TEs exhibited no or little change in their piRNA abundance (within 3^-1^ to 3^1^ folds; Figures 1B, S1D and S1E) or in their transcript abundance (Figure S1F) over the course of 7 years. Importantly, among these TEs with unchanged piRNA levels, we found that some of them exhibited remarkable changes in the pattern of piRNA production over 7 years, while others did not. For example, TE1_bm_228 (TE228) maintained the same piRNA production pattern—a sense piRNA with its 5’ end at position 7,245 and a 10-nt overlapping antisense piRNA (the ping-pong partner of the sense piRNA) at position 7,254—between Naive13 (Figure 1C, top) and Naive20 (Figure 1C, bottom). In contrast, TE1_bm_159 (TE159) had a sense piRNA at position 1,232 and its antisense piRNA partner at position 1,241 in Naive13 (Figure 1D, top), but these piRNAs decreased, and instead, a new sense piRNA at position 1,236 and its antisense piRNA partner at position 1,245 emerged in Naive20 (Figure 1D, bottom).

### The abundance of piRNAs negatively correlates with the variability of their production patterns

To comprehensively and quantitively understand the changes in piRNA production patterns in a TE-wide manner, we defined a score called the divergence score (Dscore), which represents the overall difference between two libraries in the piRNA-producing positions for each TE (Figure S2A). For example, when a TE produces piRNAs from exactly the same sites in both libraries, the Dscore of that TE is 0 (Figure S2A, Condition 1). In contrast, if a TE generates piRNAs from completely different sites in the two libraries, then the Dscore is 100 (Figure S2A, Condition 3). When a TE produces 50% of piRNAs from the same site and the other piRNAs from a different site, the Dscore is 50 (Figure S2A, Condition 2). Using this approach, we determined the Dscore distributions between our small RNA libraries. Compared to those of Naive13 vs. dsRluc13 and of Naive20 vs. dsRluc20 (Figure 2A), the distribution of Dscores was significantly higher in Naive13 vs. Naive20 (Figure 2B), indicating greater changes in piRNA production patterns. When we divided the TEs into five bins based on the Dscores of Native13 vs. Native20, the TEs categorized in each bin exhibited a consistent Dscore distribution even in the comparison of dsRluc13 vs. dsRluc20 (Figure S2B). These results suggest that the observed Dscore distributions are due to changes in piRNA production patterns over the course of seven years, rather than random variability in the library preparation process.

**Figure 2.**
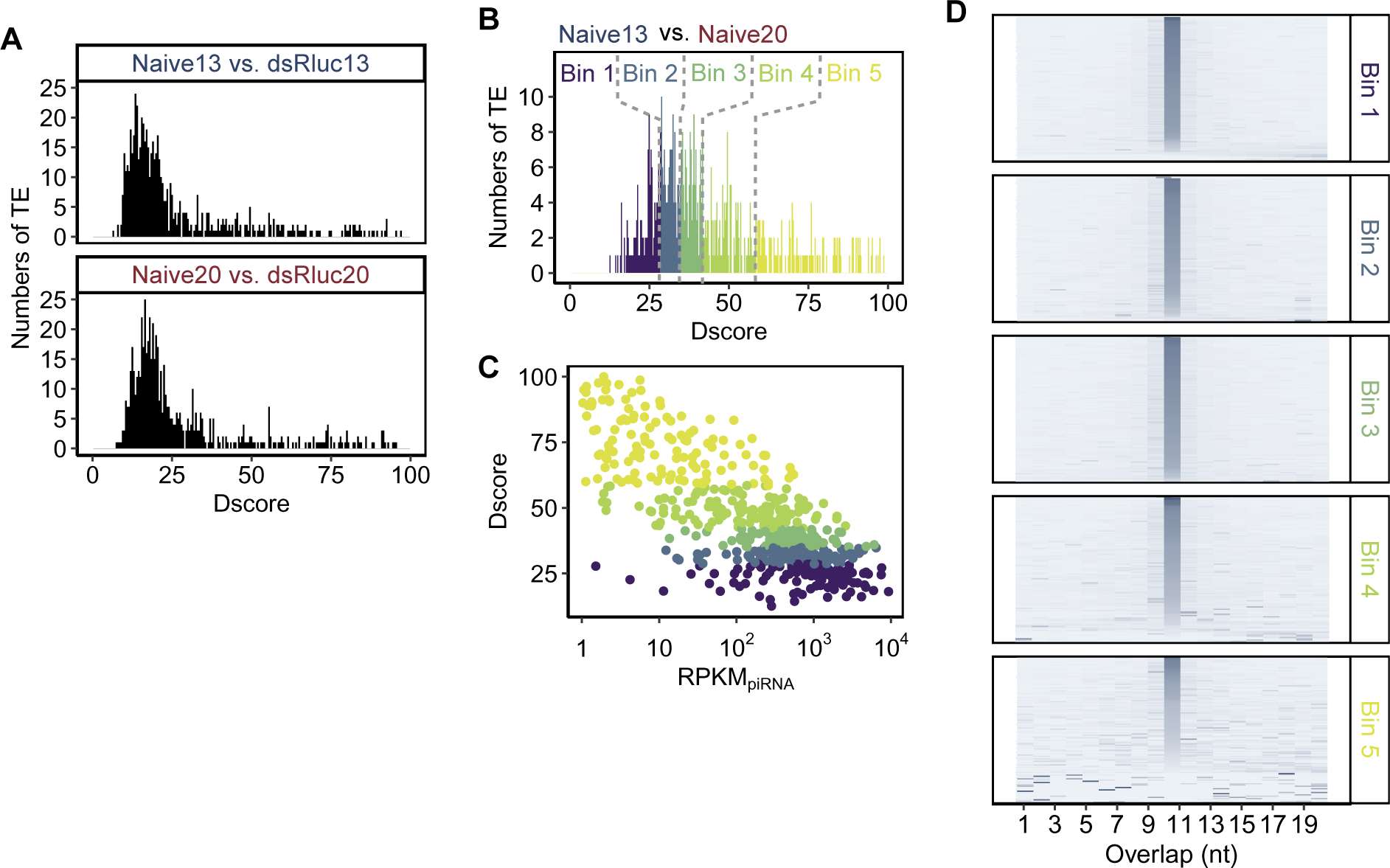
The abundance of piRNAs negatively correlates with the variability of their production patterns. (A) Histograms of Dscores between Naive13 and dsRluc13 (top) and between Naive20 and dsRluc20 (bottom) for 1,811 well-defined TEs^27^. The mean values of Dscore were 27.4 and 29.59, respectively. (B) Histogram of Dscores between Naive13 and Naive20, indicating big changes in the pattern of piRNA production. TEs were divided into five bins according to their Dscores. The mean Dscor Naive13 and Naive20 was 44.7. (C) Scatter plot showing a negative correlation between piRNA abundance (RPKM) and Dscore. Total reads mapped to 1,811 TEs^27^ were used for normalization. The x-axis represents the base-10 logarithm of RPKM. (D) Heatmaps showing the distribution of 1–20 nt overlaps of piRNAs for each TE across five bins. Enrichment of the 10-nt overlap, indicative of ping-pong-mediated biogenesis, was found in all bins.

To understand why some piRNAs changed their production sites while others did not over time, we compared the Dscore and the abundance of piRNAs for each TE. Strikingly, we observed a significant negative correlation between them with a correlation coefficient (r) of 0.41 (p < 0.01) (Figure 2C). This finding indicates that TEs with less abundant piRNAs are more prone to changes in their piRNA production patterns.

piRNAs are generally amplified by the ping-pong pathway, which produces pairs of sense and antisense piRNAs with 10-nt overlaps. To determine whether the degree of this ping-pong amplification contributes to the changeability of the piRNA-producing sites, we analyzed the enrichment for 1-to 20-nt overlaps of piRNAs for each TE in the five bins categorized based on the Dscore of Native13 vs. Native20 (Figure 2B). TEs in all five bins showed significant enrichment for the 10-nt overlaps, indicative of the ping-pong amplification of piRNAs (Figure 2D). Thus, the variability of piRNA production patterns simply depends on the abundance of piRNAs, not on how they are produced.

### The abundances of piRNAs and their precursor transcripts affect the variability of piRNA production patterns

In addition to canonical biogenesis pathways, piRNAs can also be produced from apparently random sites on TE RNAs as well as on mRNAs, presumably via non-specific degradation, at a very low frequency in a quantity-dependent manner^28,29^. Although rare, the emergence of such random piRNAs could in theory initiate a new ping-pong amplification site and may compete with a pre-existing ping-pong site(s) in the neighboring region, which may eventually lead to changes in the piRNA production patterns. To investigate how much this competition impacts piRNA production patterns, we used mathematical modeling, similar to the approach used in population genetics to study genetic drift^30,31^. In our model (Figure 3A), the red arrow and dots represent newly emerged piRNAs, while the blue arrows and dots represent pre-existing piRNAs in close proximity. The number of dots indicates the total number of piRNAs producible from this hypothetical region (N_piRNA_). In the initial state, there is always one red dot in the population. In each round, every piRNA tries to guide the production of its responder piRNA of the same color via the ping-pong pathway, with an equal chance (1/N_piRNA_). Therefore, the probability of winning the competition for a piRNA is proportional to the current population ratio between the piRNA(s) of the same color and those of the other color. If it wins, it can produce the responder piRNA of the same color; if it loses, then it is replaced with the responder piRNA of the other color. One trial continues through multiple rounds of competition until all the piRNAs in the population collapse into the same color, either newly emerged or pre-existing piRNAs. To avoid endless computations, the maximum number of rounds was set at 10,000. We ran a large number of trials (Ω_trials_) to observe the color collapse and counted the number of occurrences of successful color replacement by red dots (n_replacement_). Then, we determined the probability of red replacement, i.e., replacement of the piRNA production site by the emergence of new piRNAs.

**Figure 3.**
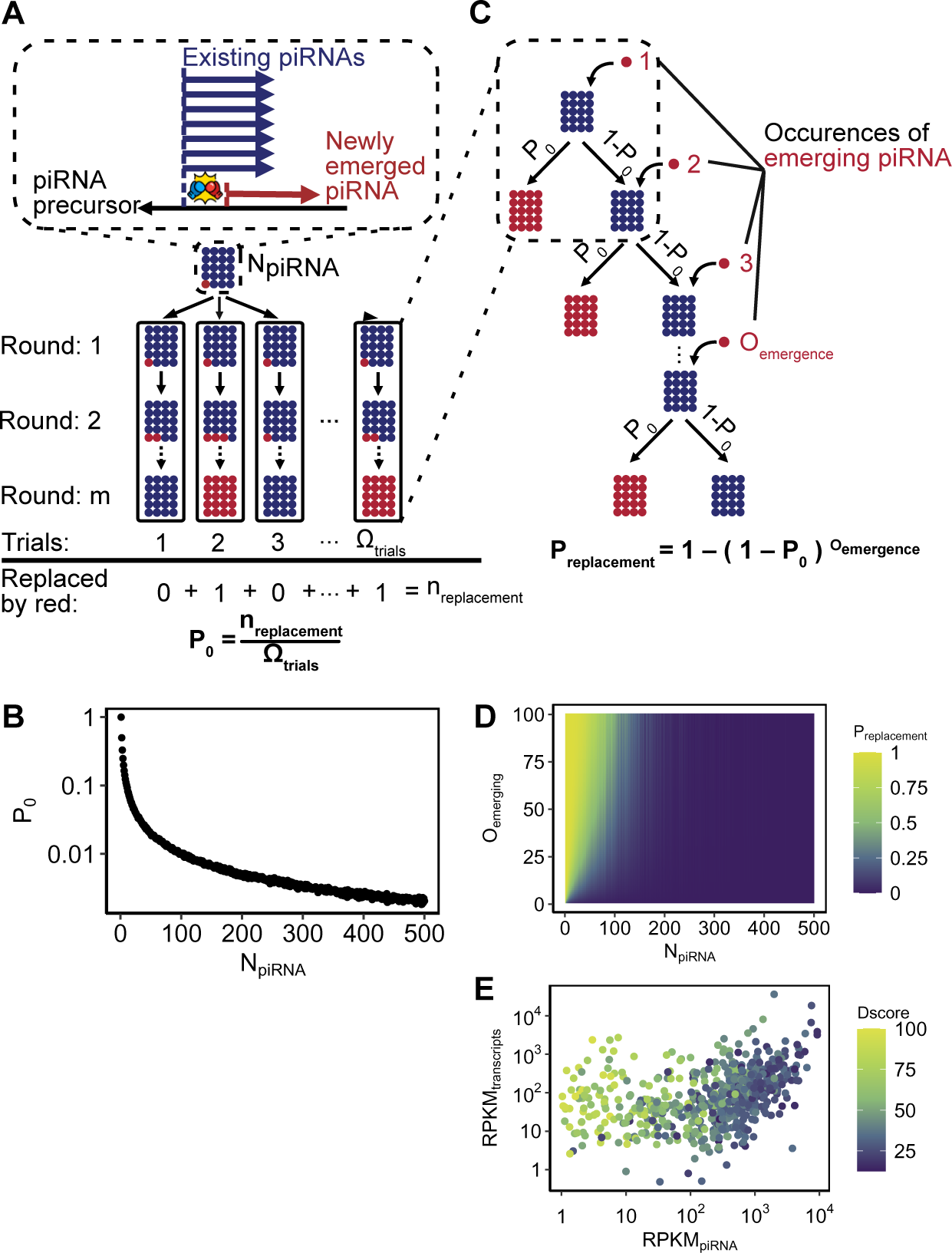
The abundances of piRNAs and their precursor transcripts affect the variability of piRNA production patterns. (A) Scheme for a mathematical modeling method starting with one newly emerged piRNA. The red dots and the arrow represent newly emerged piRNAs, while the blue dots and arrows represent pre-existing piRNAs. N_piRNA_indicates the total number of piRNAs producible from a hypothetical region. The probability of red replacement (P_0_) was calculated by dividing the number of trials when red piRNAs occupy N_piRNA_ (n_replacement_) by the total number of trials (Ω_trials_). (B) Scatter plot illustrating the relationship between N_piRNA_ (x-axis) and P_0_ (y-axis). (C) Scheme for the mathematical model that considers multiple emergences of a red piRNA after each failure of red replacement with P_0_ in (A). O_emergence_ indicates the number of red piRNA emergence. As O_emergence_ increasing, the probability that red replaces blue at least once (P_replacement_) is calculated. The rest of the model was the same as in (A). (D) Heatmap of the results obtained from the analysis of (C). The color scale represents P_replacement_, with warmer colors indicating higher probabilities and cooler colors indicating lower probabilities. (E) Heat-mapped scatter plot showing the empirical relationships among the piRNA abundance, the transcript abundance, and the Dscore between Naive13 and Naive20 for each TE. Each point represents one TE, colored according to the Dscore value ranging from cool color (low Dscore) to warm color (high Dscore). The x-axis and y-axis represent the base-10 logarithm of RPKM.

As shown in Figure 3B, our modeling indicates that the probability of replacement at the piRNA production site (P_0_; y-axis) is higher when the total number of piRNAs (N_piRNA_; x-axis) is smaller, agreeing well with our experimental observation that TEs with less abundant piRNAs are more prone to changes in piRNA production patterns (Figure 2C).

In the above modeling, we always started the simulation with one red dot in the initial state and the rest was a simple dichotomy-type competition between the two colors. In reality, however, new piRNAs can randomly (albeit infrequently) emerge multiple times, as long as their precursor molecules are available. We therefore added an assumption that, after each failure of red replacement (i.e., occupation by blue dots), one new red dot can newly emerge in the population, in addition to the red dot that was always included in the initial state (Figure 3C). The number of this new emergence of red dots (O_emergence_) depends on the availability (abundance) of their precursor molecules, given that the emergence of new piRNAs is a random event likely via non-specific degradation of precursors. As shown in Figure 3D, we observed not only that N_piRNA_ (the total number of piRNAs; x-axis) negatively correlate with the probability of replacement in the piRNA production pattern (P_replacement_; now shown in a color gradation) but also that O_emergence_ (the number of red dot emerged, which depends on the abundance of precursors; y-axis) positively correlate with the probability of replacement, especially when N_piRNA_ is small. In other words, our mathematical modeling results suggest that TEs with abundant precursors and rare piRNAs (i.e., inefficient piRNA biogenesis) are more changeable in their piRNA production patterns, whereas TEs with rare precursors and abundant piRNAs (i.e., efficient piRNA biogenesis) are more stable in their piRNA production patterns. Importantly, this trend agrees well with our experimental data regarding the relationships among piRNA abundance, transcript abundance, and Dscore (changes in piRNA production pattern) for each TE over the course of 7 years of cell passage (Figure 3E). These results highlight a simple rule behind the changeability of piRNA production patterns. When piRNA production efficiency is high, there are not many extra precursors remaining. Consequently, new piRNAs emerge less frequently, and the pattern of piRNAs is less likely to change. On the other hand, when piRNA production efficiency is low, a large number of extra precursors remain, leading to a high likelihood of new piRNAs emerging and a greater chance of piRNA pattern changes.

### piRNAs can autonomously avoid mismatches for efficient biogenesis

What is the advantage for the piRNA system to have changeable piRNA production patterns? An ongoing evolutionary arms race exists between TEs and the piRNA system. For TEs, sequence mutation is the simplest strategy to evade piRNA-mediated silencing. On the other hand, the piRNA system has more relaxed rules for target recognition and cleavage than the siRNA system, and thus it is thought to be better equipped to silence rapidly diverging TEs^20^. In general, piRNA-mediated target cleavage tolerates mismatches essentially at any position, but the base pairing status in the 5’ region (positions 2–10) of piRNAs is relatively more crucial for cleavage compared to other regions in vivo^20^. Given that each cycle in the ping-pong amplification of piRNAs relies on target cleavage, the cumulative impact of mismatches could become substantial over time. Therefore, it would be optimal for the piRNA system to avoid mismatches in the 5’ region for better silencing. Assuming a simplified scenario where piRNAs are first supplied from a piRNA cluster and then amplified by cleaving TE transcripts from other genomic loci, it must be a challenging task to precisely modify the genomic sequence of the piRNA cluster at specific positions to avoid mismatches with diverging TEs. A more effective strategy would be to adjust the piRNA-producing positions to match consensus sequences between the piRNA cluster and diverging TE copies, thereby minimizing mismatches especially in the 5’ region. However, it is highly unlikely that a “top-down” commander-like factor exists that can scrutinize all TE sequences and direct the biogenesis factors to produce piRNAs at specific positions to minimize mismatches in the 5’ region. We propose that piRNAs can overcome this challenge through a process of “self-organization,” wherein they autonomously adapt their production sites to avoid deleterious mismatches. This can be achieved by a simple mechanism in which mutations within the 5’ region of existing piRNAs decrease the efficiency of target cleavage and ping-pong amplification, thereby leaving more precursor transcripts available for the emergence of new piRNAs in close proximity that could potentially have a better base-pairing status by random chance. On the other hand, when ping-pong amplification is highly efficient with good base-pairing, most precursor transcripts are consumed by existing piRNAs for cleavage, restricting the emergence of new neighboring piRNAs and thus leading to stable piRNA production patterns. In this way, piRNA production patterns can be autonomously optimized, maintained, and re-optimized, as a system, to avoid deleterious mutations and reinforce the silencing of continuously mutating TEs.

To explore this idea, we analyzed the positional distribution of mismatches to the consensus sequences within the 5’ (positions 2–9), middle (11–18), and 3’ (19–26) regions of piRNAs in each bin of TEs (Figure 4A). We found that piRNAs in bin 1 TEs, characterized by efficient piRNA biogenesis and stable production patterns, tend to have mismatches predominantly in the middle and 3’ regions, avoiding more harmful mismatches in the 5’ region (Figure 4B). In contrast, piRNAs in bin 5 TEs, which are characterized by inefficient biogenesis and changeable production patterns, did not exhibit this avoidance of 5’ mismatches; instead, mismatches were more evenly distributed across all regions of bin 5 piRNAs (Figure 4B). Therefore, piRNAs in bin 1 TEs appear to be more optimized than those in bin 5 TEs by avoiding deleterious mismatches in their 5’ regions, suggesting that the stability of piRNA production patterns is determined at the level of individual TEs in cultured BmN4 cells. Notably, this trend was observed for both Naive13 and Naive20 (Figure 4B), suggesting that the optimization of piRNA production patterns has already reached saturation in BmN4 cells or that at least 7 years of routine cell passage under laboratory conditions is not sufficient to promote further optimization.

**Figure 4.**
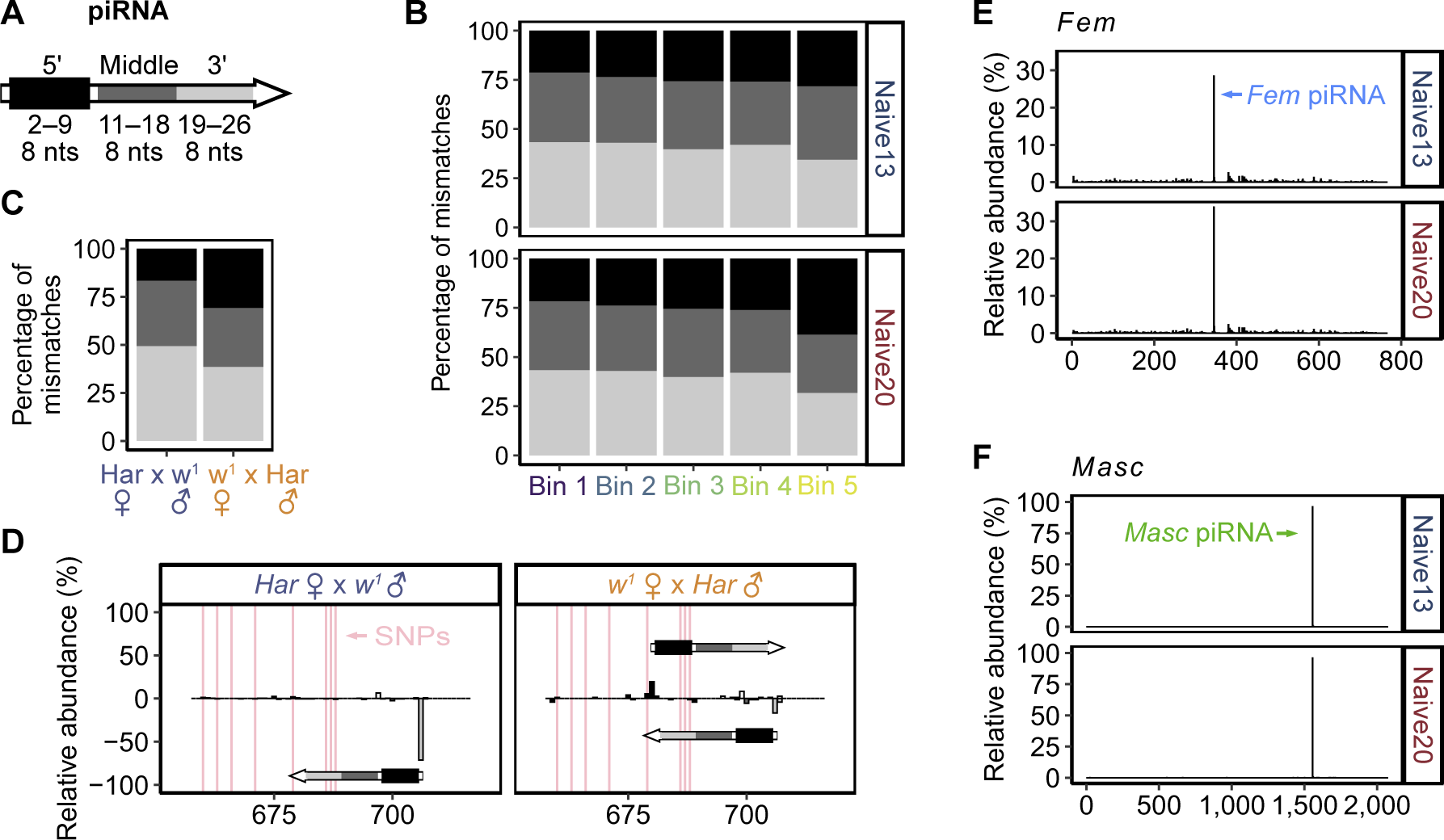
piRNAs can autonomously avoid mismatches for efficient biogenesis. (A) Scheme for the definition of piRNA regions: 5’ region (nt positions 2–9 from the piRNA 5’ end, black), middle region (nt positions 11–18, grey), and 3’ region (nt positions 19–26, light grey). (B) Bar plot illustrating the distribution of mismatches across the three piRNA regions. The x-axis represents the five bins according to the Dscores in Figure 2B, while the y-axis represents the percentage of mismatch in each piRNA region defined in (A). (C) The proportion of mismatches in the three regions of all piRNAs that map to the entire *P* element. These piRNAs were originated from 21-day-old dysgenic hybrids (*w^1^* female x *Har* male, right) and reciprocal hybrids (*Har* female x *w^1^* male, left). (D) Bar plot illustrating the distribution of the 5’ ends of piRNAs derived from the dysgenic hybrids (right) and reciprocal hybrids (left) that mapped to a specific region of *P* element. Total reads in this region were used for normalization. Representative piRNA, which have the SNP positions within the three regions defined by (A) are illustrated. The pink vertical lines represent the positions of SNPs (660, 663, 666, 671, 679, 686, 687 and 688). (E and F) Bar plot illustrating the distribution of the 5’ ends of piRNAs that mapped to *Fem* (E) and *Masc* (F), separately showing one main peak at position 345 from *Fem* and one main peak at position 1,556 from *Masc* in both Naive13 (top) and Naive20 (bottom). Total reads mapped to *Fem* and *Masc* were used for normalization, respectively.

Therefore, we decided to shift our focus to the phenomena of *de novo* production of piRNAs previously reported in *Drosophila*^22,32,33^. In the case of P-M hybrid dysgenesis in *Drosophila*, when a male with a TE called *P* element (*Harwich* strain [*Har*]) mates with a female lacking a *P* element (*w^1^*), the resulting female offspring are almost infertile at a young age due to the lack of maternally inherited *P* element-targeting piRNAs^22,34,35^. However, the fertility of these dysgenic hybrid females improves as they age, because *P* element piRNAs are produced *de novo* from paternally inherited clusters over time^22^. These *de novo* piRNAs are produced from scratch within a single generation, without the potential benefit of trans-generational inheritance and re-amplification. Therefore, they are expected to have a sequence repertoire that is not as fully optimized as that of maternally inherited piRNAs in the reciprocal non-dysgenic hybrids. To explore this idea, we analyzed the sequences of *P* element-targeting piRNAs in dysgenic hybrids (*w^1^* female × *Har* male) and reciprocal hybrids (*Har* female ×*w^1^* male). Specifically, we examined their positional relationships with the sites of sequence variations (SNPs) on the active *P* element, which could cause mismatches against piRNAs. As expected, we found that piRNAs in reciprocal hybrids tend to avoid mismatches in the 5’ region, suggesting optimized piRNA production patterns (Figure 4C, left). In contrast, piRNAs in dysgenic hybrids exhibited mismatches rather evenly across the 5’, middle, and 3’ regions (Figure 4C, right). For example, piRNAs in dysgenic hybrids are promiscuously produced from a *P* element segment that contains eight SNPs (660, 663, 666, 671, 679, 686, 687, and 688), with one of the major piRNAs even having three SNPs in its 5’ region (Figure 4D, right). However, piRNAs in reciprocal hybrids were primarily produced from a single site with SNPs located in the non-detrimental 3’ region (Figure 4D, left). These results suggest that the sequence repertoire of *do novo* piRNAs is initially immature, but it can presumably be optimized over generations through maternal deposition and re-amplification.

## DISCUSSION

In summary, our study illustrates how competition between neighboring ping-pong sites autonomously shapes the repertoire and biogenesis efficiency of piRNAs (Figure S3). We showed that piRNA-producing sites are highly changeable when their biogenesis efficiency is low. This mechanism is similar to genetic drift in population genetics, where the change in allele frequencies through random sampling is more pronounced when the population size is small^30,31^. Moreover, the emergence of new piRNAs is reminiscent of gene flow, the transfer of genetic material from one population to another, in population genetics^30,31^. However, there is a crucial difference: In population genetics, there is no general relationship between the frequency of gene flow and the genetic fitness of a population^30,31^. However, in the piRNA system, the frequency of new piRNA emergence depends on the availability of unconsumed precursor transcripts, which inversely correlates with the efficiency of piRNA production (corresponding to genetic fitness) (Figures 3D and 3E). As such, the piRNA system can autonomously explore more optimized production patterns when their biogenesis efficiency is low. Conversely, when the biogenesis efficiency is high, optimized piRNA production patterns can be automatically stabilized. This unique feature enables a flexible balance between dynamic adaptation and stable maintenance of piRNA production patterns, ensuring effective defense against diverse TEs. Crucially, all of this is made possible by simple competition between adjacent ping-pong sites, because the ping-pong mechanism intrinsically couples the cleavage of target TEs and the amplification of new piRNAs. Our study not only provides the biological reason for the vastly complex ping-pong pathway of piRNAs but also highlights a universal principle in biology: the power of competition and self-amplification to drive autonomous optimization.

In general, ping-pong amplification of transposon-derived piRNAs occurs between sense and antisense transcripts that have high complementarity over long ranges. However, some specialized piRNAs are produced through heterotypic ping-pong amplification between two distinct transcripts; these piRNAs originate from separate genomic loci and exhibit complementarity only within a short region (Figure S4). For example, a female-specific, sex-determining piRNA is produced from a precursor transcript called *Feminizer* (*Fem*) in silkworms^36^. *Fem* piRNA cleaves a complementary sequence in the mRNA of a gene named *Masculinizer* (*Masc*), the protein product of which governs masculinization; silencing of *Masc* by *Fem* piRNA is essential for feminization. The *Fem* transcript and *Masc* mRNA exhibit complementarity solely at the *Fem* and *Masc* piRNA target site, therefore even a single nucleotide shift in the *Fem* piRNA-producing site would prevent its biogenesis via heterotypic ping-pong. Indeed, *Fem* and *Masc* piRNAs showed the identical production patterns between Native13 and Naive20 in our analysis (Figures 4E and 4F). Thus, these sex-determining piRNAs deviate from the intrinsic flexibility and adaptivity of piRNA production sites, thereby ensuring their critical biological roles beyond TE silencing.

The piRNA system is a highly conserved mechanism across the animal kingdom, with many species utilizing the ping-pong pathway for piRNA biogenesis^5,37-40^. However, recent findings that some *Drosophila* species lack the ping-pong pathway highlight the potential for maintaining a functional piRNA system without relying on this mechanism^41^. The ping-pong pathway is vastly complex and presumably costly; it requires numerous protein factors including many ATP-dependent helicases and precise regulations across multiple subcellular compartments, which may pose an evolutionary trade-off. In some situations, simple expansion of the piRNA sequence repertoire by the phased pathway may be sufficient to silence TEs. On the other hand, the widespread conservation of the ping-pong pathway suggests a selective advantage in autonomously and flexibly optimizing the piRNA repertoire in response to constant changes of TEs. Further studies are warranted to elucidate the interplay between these two piRNA biogenesis pathways in maintaining the genome integrity over many generations under the threats of rapidly diverging or newly acquired TEs.

## ACKNOWLEDGMENTS

We thank the members of Tomari laboratory for critical comments on the manuscript. This work was supported in part by Grant-in-Aid for Scientific Research (S) (18H05271 to Y.T.), Grant-in-Aid for Scientific Research (A) (23H00364 to Y.T.), Grant-in-Aid for Early-Career Scientists (22K15082 to K.S.).

## AUTHOR CONTRIBUTIONS

J.Y., K.S. and Y.T. conceived and designed the experiments and wrote the manuscript. N.I. prepared the small RNA libraries. J.Y. performed the bioinformatics analyses. K.S. and Y.T. supervised the research. All the authors contributed to the discussion of the results and the approval of the manuscript.

## METHODS AND MATERIALS

### Cell lines

All BmN4 cell lines were compared in this study: Naive13 (maintained in our laboratory and stocked at –80°C from 2013 to 2020), Naive20 (originated from naive13 and continuously cultured for 7 years), dsRluc13 and dsRluc20 (prepared by RNAi as described previously^15^).

### Small RNA library preparation

Small RNA libraries were constructed according to the Zamore lab’s open protocol (https://www.dropbox.com/s/r5d7aj3hhyaborq/) with several modifications. The 3′ adapter was conjugated with an amino CA linker instead of dCC at the 3′ end (GeneDesign) and then adenylated at the 5′ end with a 5′ DNA adenylation kit (NEB). Four random nucleotides were added in the 3′ and 5′ adapters [(5′ -rAppNNNNTGGAATTCTCGGGTGCCAAGG/amino CA linker-3′) and (5′ -GUUCAGAGUUCUACAGUCCGACGAUCNNNN-3′)] to decrease ligation bias. The adaptor ligation was performed with the addition of 20% PEG-8000. Small RNA libraries were sequenced using the Illumina HiSeq 4000 platform to obtain single end reads.

### Sequence analysis of small RNA library

The small RNA reads generated by Illumina HiSeq 4000 sequencing had a length of 36 nucleotides (nt). The 3′-adaptor sequences were identified and removed by cutadapt^42^, allowing for up to two mismatches. Reads shorter than 26 nt or longer than 32 nt were excluded, resulting in reads ranging from 26 to 32 nt. To map the small RNAs, the 1,811 *Bombyx* TEs^27^ and sequences of *Fem* and *Masc^36^* were used. The mapping process was carried out using the Bowtie aligner^43^, allowing for up to three mismatches between the reads and the reference sequences. The alignment results were used to calculate the mapping rate of each library, with mapping rates against the 1,811 TEs^27^ utilized for normalization. The resulting alignment files in SAM format were converted to BAM format using SAMtools^44^. Subsequently, BEDTools^45^ was used to convert the BAM files into bed format. From the bed files, the 5’-end positions of each piRNA were obtained using R programs.

Four distinct samples were used for the calculation of Dscore: Naive13, Naive20, dsRluc13, and dsRluc20. For each sample, piRNA data, encompassing reads per million (RPM) for total reads, was collected. Subsequently, the reads per kilobase per million mapped reads (RPKM) for each TE element were computed. The RPKM values were normalized by each TE’s total RPKM.

#### Calculation of Dscore

Dscore was calculated for each pair of sample sets (e.g., Naive13 vs dsRluc13). The computation involved determining the absolute difference of RPKM values for corresponding positions between the two samples and summing these differences. For each individual position, the RPKM values were extracted from both samples. Following this, the difference between these two values was calculated. Importantly, this is an “absolute difference,” meaning that regardless of which number is greater, by subtracting the smaller value from the larger, a positive number is always obtained. This ensures that influences from diverging positive and negative differences are mitigated. Lastly, to obtain the total difference for each TE between the two biological samples, all these discrepancies at these respective positions within the TE were summed up. Only TEs with a length longer than 1,000 base pairs were considered.

#### Calculation of Ping-Pong Signature in piRNA Sequences

Ping-pong signature was calculated by the amounts of plus and minus strand RPKM values at every position along the piRNA sequences, using a sliding window approach.

#### The positional distribution of mismatches within piRNAs

piRNA sequences were aligned with the consensus sequence of piRNA at each position associated with the identified TE bins. The consensus sequence was obtained by using “DECIPHER” in R (https://www.deciphergenomics.org) to extract the most frequent sequence at each position. The number and types of mismatches within the three regions of piRNAs: 5’ (positions 2–9), Middle (positions 11–18), and 3’ (positions 19–26), were determined using R programs.

#### The positional analysis of *P* element

The *P* element sequence was extracted from AB331393.1 of GenBank at NCBI, and the region from base 246 to base 3,152 corresponding to the *P* element was used. Data from a previous study (SRP007937) was downloaded from SRA, and for the small RNA sequence, the adaptor was removed by cutadapt^42^ and mapped to the *P* element by Bowtie^43^, allowing up to 3 nucleotide mismatches. For genomic DNA-seq, mismatches of up to 3 nucleotides were allowed by Bowtie and mapped to the *P* element. From the genomic DNA-sequence results, we reconstructed the correct sequence of the *P* element by extracting the parts of the sequence that differed from the reference using our R script. SNPs were defined as those regions where more than 5 reads of the sequence differed from the consensus. The distribution of the 5’ ends of small RNAs mapped to these regions was visualized by R.

### RNA library preparation

Total RNAs were isolated from BmN4 cells using TRIzol reagent (Invitrogen) referencing to the manufacturer’s protocol. Subsequently, RNA samples were separately entrusted to QB3-Berkeley Genomics core labs (Naive13) and BGI, Hong Kong (Naive20) for polyA selection, the preparation of RNA libraries and sequencing.

### Sequence analysis of RNA library

The RNAs were mapped to the 1,811 *Bombyx* TEs^27^. The mapping process of RNAs was carried out using the hisat2^46^. The resulting SAM files were converted to BAM format by SAMtools^44^, and then the BAM files were converted to bed format using BEDTools^45^. From the bed, the RPKM of RNA from each TE was calculated using R programs.

### Simulation on the dynamics of piRNA production and competition

The computational model references the Moran process^30^, and our model is based on the ping-pong pathway, a mechanism for generating secondary piRNAs from primary transcripts. We assume that the ping-pong pathway operates in a stochastic and competitive manner, so that each piRNA has an equal chance of being replaced by another piRNA from a different source.

#### Probability of red replacement (P_0_)

Simulation for the total number of piRNAs (N_piRNA_) was defined ranging from 1 to 500. Each simulation consisted of a total of 100,000 trials (Ω_trials_). In each trial, there were m rounds of iterations, with each round starting with one newly emerged piRNA in the N_piRNA_ as the initial state. The initial round of each trial involved randomly sampling successes (sampling newly emerged piRNAs) based on a given probability (1/N_piRNA_). The count of successes was then calculated as a proportion of the total sampling attempts (N_piRNA_), serving as the random sampling probability for the next round. This process continues until the probability collapses to 0 (complete disappearance of newly emerged piRNAs) or 1 (newly emerged piRNAs occupy the entire N_piRNA_). The maximum value for m would not exceed 10,000 to prevent infinite iterations. Finally, the probability of red replacement (P_0_) is calculated by the proportion of times red piRNAs occupy N_piRNA_ (n_replacement_) in total trials (Ω_trails_).

#### Probability that red replaces blue at least once (P_replacement_)

Assuming the number of red piRNA emergence (O_emergce_) has a range from 1 to 100, calculate the P_replacement_ based on the following formula: P_replacement_ = 1 - (1 - P_0_) O_emergence_.

### Data access

Deep sequencing data obtained in this study are accessible through the provided accession number DRA017494 (DDBJ). The *Drosophila P* element piRNA and DNA sequences (SRP007937) were reported previously^26^.

### Declaration of generative AI and AI-assisted technologies in the writing process

During the preparation of this work the authors used ChatGPT in order to improve readability and language. After using this tool/service, the authors reviewed and edited the content as needed and take full responsibility for the content of the publication.

### Code availability

All code used in this study is available on GitHub at https://github.com/YJ-July/auto-shaping.git

