## Supplementary material for "Autonomous Shaping of the piRNA Sequence Repertoire by Competition between Adjacent Ping-Pong Sites": Fig. S1-S4

### Figure S1

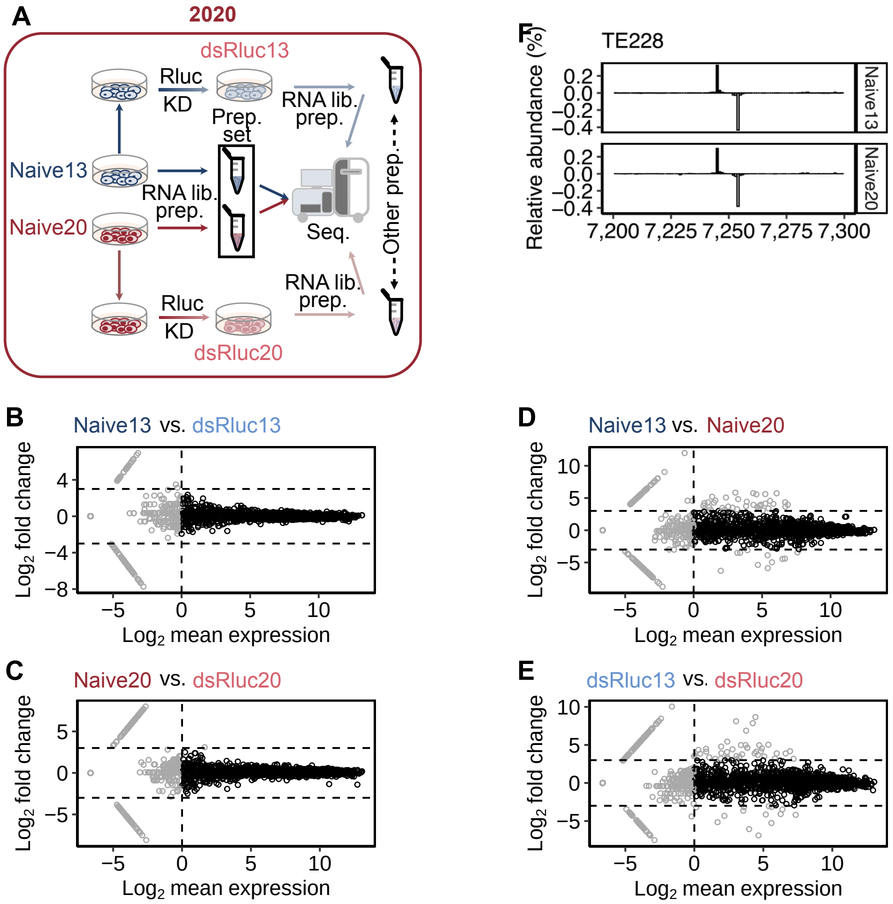


**Figure S1. TEs showed little changes both in their piRNA levels and transcript levels, related to Figure 1.**

(A) Schematic description of the dsRluc13 and dsRluc20 libraries. The dsRluc13 and dsRluc20 libraries were prepared from Naive13 and Naive20 cells transfected with dsRNAs against Renilla luciferase as mock RNAi, as described previously[15].

(B–E) MA plots depicting differential piRNA expression analysis for each TE. Each point represents an individual TE. TEs showing no differential piRNA expression are colored in black. Only TEs with mean values exceeding 1 RPKM are shown in the plots.

(F) MA plot depicting the differential TE transcript expression analysis. Each point represents an individual TE. TEs with no differential piRNA expression, shown in black in (D), are highlighted with black in the plot.

#### Figure S2

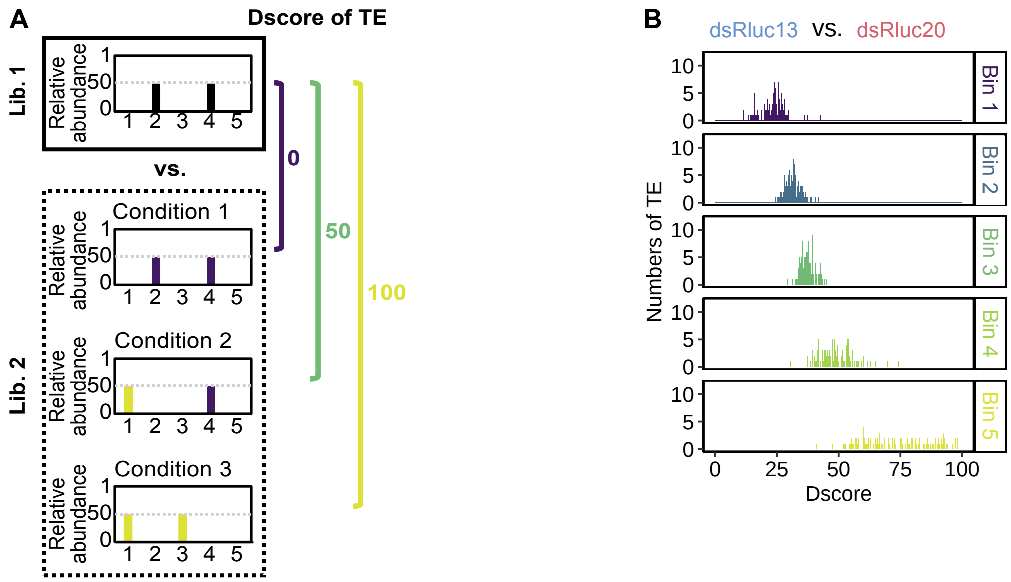


**Figure S2. Dscores reflects the changes in the piRNA production patterns, related to Figure 2.**

(A) Scheme for the calculation of the divergence score (Dscore) for a TE, which represents the degree of changes in the piRNA production patterns. For each TE, the ratio of piRNAs at each piRNA production site was first calculated. The Dscore was then calculated based on the sum of the difference in the ratio for each site between two libraries.

(B) Analysis of TEs based on Dscores between dsRluc13 and dsRluc20. TEs were divided into five bins according to the Dscores between Naive13 and Naive20.

#### Figure S3

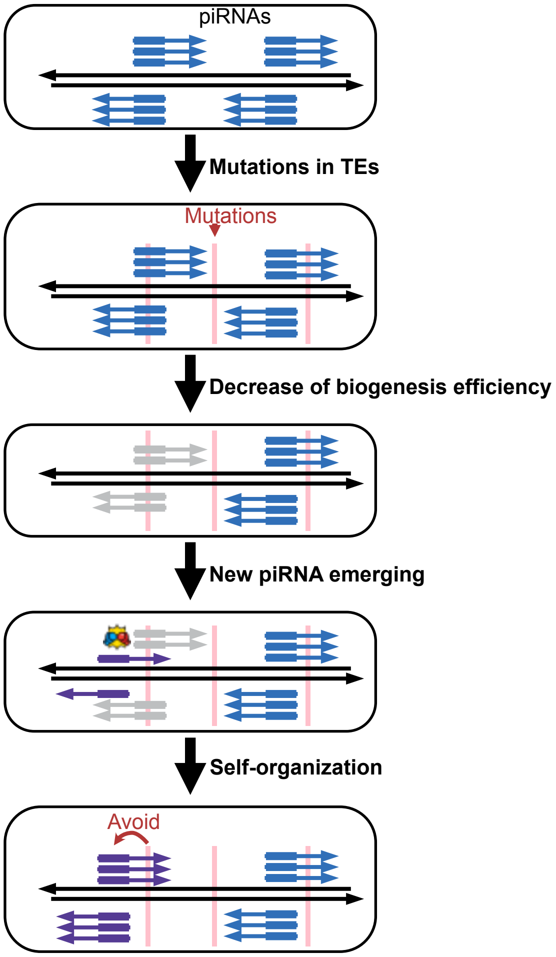


**Figure S3. Competition between neighboring piRNA-producing sites autonomously shapes the repertoire of piRNAs, related to Figure 4.**

Model for the piRNA self-organization mechanism. The occurrence of mismatches (indicated by the red shading) within a stable piRNA generation system (blue), particularly when they appear in the 5' region (grey), which is critical for the efficiency of piRNA production, leads to the impaired production of piRNAs at those positions. At the same time, new piRNAs (purple) randomly emerge at neighboring locations. Through competition, these new piRNAs with better cleavage efficiency can replace the preceding piRNAs through self-organization, thus avoiding critical 5' regions via mismatch positions.

#### Figure S4

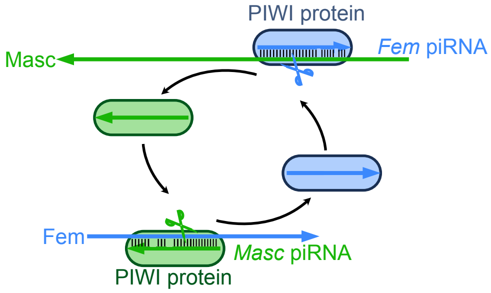


**Figure S4. The *Fem* transcript and Masc mRNA exhibit complementarity solely at the *Fem* and *Masc* piRNA target site, related to Figure 4.**

Scheme for the ping-pong cycle between *Fem* piRNA and *Masc* piRNA. Short arrows represent *Fem* piRNA (sky blue) and *Masc* piRNA (green) targeting at *Masc* mRNA (long green arrow) and *Fem* mRNA (long sky-blue arrow) at position 1,556–1,584 and position 345–373, respectively.
